# The effects of microcystin-LR in *Oryza sativa* root cells: F-actin as a new target of cyanobacterial toxicity

**DOI:** 10.1101/2020.02.17.952218

**Authors:** D. Pappas, S. Gkelis, E. Panteris

## Abstract

- Microcystins are toxins produced by cyanobacteria, notorious for negatively affecting a wide range of living organisms, among which several plant species. Although microtubules are a well-established target of microcystin toxicity, its effect on filamentous actin (F-actin) in plant cells has not been studied yet.
- The effects of microcystin-LR (MC-LR) and the extract of a microcystin-producing freshwater cyanobacterial strain (*Microcystis flos-aquae* TAU-MAC 1510) on the cytoskeleton (F-actin and microtubules) of *Oryza sativa* (rice) root cells, were studied by light, confocal, and transmission electron microscopy. Considering the role of F-actin in endomembrane system distribution, the endoplasmic reticulum and the Golgi apparatus in extract-treated cells were also examined.
- F-actin in both MC-LR- and extract-treated meristematic and differentiating root cells exhibited time-dependent alterations, ranging from disorientation and bundling to the formation of ring-like structures, eventually resulting to a collapse of the F-actin network at longer treatments. Disorganization and eventual depolymerization of microtubules, as well as abnormal chromatin condensation were observed following treatment with the extract, effects which could be attributed to microcystins and other bioactive compounds. Moreover, cell cycle progression was inhibited in extract-treated roots, specifically affecting the mitotic events. As a consequence of F-actin network disorganization, endoplasmic reticulum elements appeared stacked and diminished, while Golgi dictyosomes appeared aggregated.
- These results support that F-actin is a prominent target of MC-LR, both in pure form and as an extract ingredient. Endomembrane system alterations can also be attributed to the effects of cyanobacterial bioactive compounds (including microcystins) on F-actin cytoskeleton.

## INTRODUCTION

Cyanobacteria are photosynthetic, oxygenic prokaryotes, inhabiting a variety of aquatic and terrestrial environments, surviving even extreme conditions (such as high temperature or salinity etc.; Codd et al., 2017). A wide range of species are known to produce cyanotoxins, a group of chemically diverse secondary metabolites (as reviewed by Pantelić et al., 2013), proven to be harmful to higher eukaryotes, including humans (Buratti et al., 2017) and various plant species (Máthé et al., 2013; Mitrovic et al., 2004). The most common cyanotoxins found in freshwater bodies are microcystins (MCs), water-soluble monocyclic heptapeptides, exhibiting two variable amino acids (Catherine et al., 2017). The toxicity of MCs is due to their ability to inhibit the activity of protein phosphatases 1 (PP1) and 2A (PP2A) (MacKintosh et al., 1990), which are involved in the cell cycle progression (Brautigan and Shenolikar, 2018).

MCs are released into the surrounding water during cyanobacterial cell lysis (Sivonen and Jones, 1999), thus becoming eventually accessible to consumers either directly (through water supply) or indirectly, affecting crops (through irrigation water) destined for consumption. When present in irrigation water (due to naturally occurring cyanobacteria in freshwaters), MCs have been shown to bioaccumulate in cultivated plant species (Corbel et al., 2016; Drobac et al., 2017), raising serious concerns over food safety and the impacts on consumers’ health (Cao et al., 2018a).

Rice (*Oryza sativa* L.) is a crop of great commercial value worldwide (FAO, 2018). Since its cultivation is closely related to the aquatic environment, it is prone to all the dangers imposed by the presence of cyanobacteria in the surrounding water. Although many studies have focused on the accumulation and toxicity of MCs in rice (Azevedo et al., 2014; Chen et al., 2012; Liang et al., 2016), their effects on the physiology of rice cells remain unclear. A recent study has underlined the negative effects of MC-contaminated water on rice root growth (Cao et al., 2018b), as a result of mitotic disruption in dividing root cells.

Previous studies on plant cells have shown that MCs disrupt microtubule organization, leading to mitotic abnormalities, and induce chromatin hypercondensation, due to histone H3 hyperphosphorylation, while alterations in cell cycle progression have also been reported (for a review on the effects of MCs, see Máthé et al., 2013). Even though chromatin and microtubules are well-known “targets” of MCs in plant cells, the other component of the plant cytoskeleton, F-actin, has not been studied in MC-treated cells of any plant species so far. However, F-actin is important for plant growth, regulating the intracellular distribution of several organelles (Volkmann and Baluška, 1999) and driving cytoplasmic streaming, a function pivotal for plant cell viability (Shimmen and Yokota, 2004). All reports about the adverse effects of MCs on plant cytoskeleton have been based on applying purified MCs (e.g. Garda et al., 2016; Máthé et al., 2013), as purified toxins offer high precision in determining the exact concentrations. On the other hand, crude extracts of MC-producing cyanobacterial strains, known to negatively affect plant growth, are helpful in simulating the natural exposure of plants to MCs during lab experiments (Pflugmacher et al., 2007; Prieto et al., 2011). In accordance, in the present study, the effects of both purified microcystin-LR (MC-LR) and a toxic cyanobacterial extract (containing MCs) on F-actin of rice root cells were investigated. In order to further highlight the significance of any disruption of F-actin due to the toxic extract, endoplasmic reticulum and Golgi apparatus distribution, related to F-actin (Boevink et al., 1998), as well as cytoplasmic streaming (Shimmen and Yokota, 2004; Volkman and Baluška, 1999) were examined.

## MATERIALS AND METHODS

### Cyanobacterial strains, growth media and culture conditions

A strain of the TAU-MAC culture collection (Gkelis and Panou, 2016), *Microcystis flos-aquae* TAU-MAC 1510, isolated from Lake Pamvotis, Greece (Gkelis et al., 2015a), was used for experimental purposes. *Microcystis flos-aquae* TAU-MAC 1510 has been found to be toxic (Gkelis et al., 2015a), producing a range of microcystins, including MC-YR, MC-LR, [D-Asp^3^] MC-LR and MC-HilR (Gkelis et al., 2019). The strain was cultured in BG-11 medium, containing NaNO_3_ (Rippka, 1988), in a 500 mL glass Erlenmeyer flask (250 mL of medium were inoculated with a 4-5 mL inoculum, from an exponentially growing pre-culture, under aseptic conditions). The culture was grown at 24±1°C for about 30 days in a 12 h:12 h light:dark cycle at a photosynthetic photon flux density of 10 μmol m^-2^ s^-1^ using cool white fluorescent lamps.

### Biomass collection and extraction

The culture was centrifuged at 3,500 rpm for 10 min. The supernatant was discarded and the precipitate (biomass) was stored at −70°C overnight. Frozen biomass was lyophilized with an ALPHA 1-4 freeze dryer (Martin Christ, Gefriertrocknungsanlagen, Osterode am Harz, Germany), at temperature ranging from −48°C to −54°C and pressure between 0.05 and 0.02 mbar, until dry. Dry biomass was weighed (150 mg) and extracted thrice in 21 mL of 75% (v/v) methanol in glass tubes. The sample was sonicated during the first extraction step for 10 min with a Vibra-Cell VC-300 High Intensity Ultrasonic Processor (Sonics and Materials Inc., Newtown, CT, USA) and stirred for 45 min at each extraction step at room temperature. The extract was left to evaporate under aseptic conditions and the pellet was resuspended in 5 mL of double-distilled water. The aqueous extract was filtered through Whatman Polydisc TF filters (Whatman plc, Little Chalfont, UK) with a pore size of 0.2 μm.

### Plant material and exposure to MC-LR and the crude extract

Rice (*Oryza sativa* cv Axios, kindly provided by the National Cereal Institute, Thessaloniki, Greece) seeds were germinated on filter paper moistened with tap water, in the dark, at 24±1°C. MC-LR purified from *Anabaena* strains (Halinen et al., 2007), generously provided by Prof. Kaarina Sivonen (Department of Microbiology, University of Helsinki), was used for treatments. Four- to five-day-old seedlings were placed with their roots submerged in either cyanobacterial aquatic extract or an aquatic solution of MC-LR inside Eppendorf tubes for various time periods (30 min, 1, 2, 3 or 24 h) at the same conditions. Seedlings submerged in tubes with distilled water were used as control. The concentration of the MC-LR aquatic solution was set at 45 μg·mL^-1^, equal to the total concentration of microcystins contained in the *Microcystis flos-aquae* TAU-MAC 1510 strain (Gkelis et al., 2019), in order to achieve comparable results. Following exposure, root tips 2-3 mm long were cut with steel razor blades and prepared for fluorescence and transmission electron microscopy (TEM). All chemicals and reagents were purchased from Applichem (Darmstadt, Germany), Sigma-Aldrich (Taufkirchen, Germany) and Merck (Darmstadt, Germany) and all experimental procedures described below were performed at room temperature, unless otherwise stated.

### Endoplasmic reticulum and tubulin immunolabeling

Endoplasmic reticulum and tubulin immunostaining were performed as mentioned by Adamakis et al. (2016), with some modifications. In particular, root tips were fixed in 4% (w/v) paraformaldehyde (PFA) solution in PEM buffer (50 mM PIPES, 5mM EGTA, 5 mM MgSO_4_, pH 6.8) with the addition of 5% (v/v) dimethyl sulfoxide (DMSO) for 1 h. Fixed specimens were washed with PEM (3 x 10 min) and cell walls were digested with 2% (w/v) Macerozyme R-10 + 2% (w/v) cellulase Onozuka R-10 (Duchefa Biochemie, Haarlem, Netherlands) solution in PEM for 1 h. After washing with PEM (3 x 10 min), the root tips were squashed on poly-L-lysine-coated coverslips, left to dry, and the cells were extracted with a 5% (v/v) DMSO + 1% (v/v) Triton X-100 solution in phosphate-buffered saline (PBS, pH 7.2) for 1 h. For endoplasmic reticulum immunolabeling, mouse anti-HDEL antibody (2E7, Santa Cruz Biotechnology, Dallas, TX, USA), diluted 1:50 in PBS, and AlexaFluor488-anti-mouse (Invitrogen, Carlsbad, CA, USA), diluted 1:150 in PBS, were used. For tubulin immunolabeling, rat anti-α-tubulin (YOL 1/34, Serotec, Kidlington, UK) was applied as primary antibody and incubated overnight, then washed with PBS and incubated with FITC-anti-rat at 37°C for 3 h. Both antibodies were diluted 1:40 in PBS. DNA was counterstained with DAPI (0.9 mM stock solution of 4’,6-diamidino-2-phenylindole in DMSO and further diluted 1:1000 in PBS) for 5 min. After final wash with PBS, all specimens were mounted with an anti-fade medium [PBS 1: 2 glycerol (v/v) + 0.5 % (w/v) p-phenylenediamine].

### F-actin labeling with phalloidin

F-actin labeling with fluorescent phalloidin was performed according to Stavropoulou et al. (2018). In short, F-actin in root tips was pre-stabilized in 300 μM *m*-maleimidobenzoyl-*N*-hydroxysuccinimide ester in PEM, with the addition of 0.1% (v/v) Triton X-100 for 30 min in the dark. Immediately after pre-stabilization, fixation was performed with 4% (w/v) PFA in PEM + 5% (v/v) DMSO + 0.1% (v/v) Triton X-100, while DyLight 554-phalloidin (Cell Signaling Technology, Danvers, MA, USA) 1:400 was also added in the fixative for better F-actin preservation. After washing with PEM (3 × 10 min), the specimens were extracted in 5% (v/v) DMSO + 1% (v/v) Triton X-100 in PBS for 1 h and F-actin labeling was performed by incubating with DyLight 554-phalloidin diluted 1:40 in PBS + 0.1% (v/v) Triton X-100 at 37°C for 2 h in the dark. After DNA counterstaining with DAPI (as mentioned above) and washing with PBS, all specimens were mounted with anti-fade medium.

### Confocal fluorescence microscopy

Fluorescent specimens were observed with a Zeiss Observer.Z1 (Carl Zeiss AG, Munich, Germany) microscope, equipped with the LSM780 confocal laser scanning (CLSM) module, with the appropriate filters for each fluorophore. Imaging was achieved with ZEN2011 software according to the manufacturer’s instructions.

### Fluorescence intensity measurements

Fluorescence intensity measurements for F-actin were performed in maximum intensity projections of serial CLSM sections of root tips (in the meristematic and differentiation zone), treated with 45 μg·mL^-1^ MC-LR or the cyanobacterial extract for various time periods (30 min, 1, 2 or 24 h), using ImageJ (https://imagej.net/Fiji), according to Adamakis et al. (2014) and Mylona et al. (2020). The corrected total cell fluorescence (CTCF; Gavet and Pines, 2010) was calculated with the formula: CTCF = Integrated Density – (Area of selected cell X Mean fluorescence of background readings). Thirty individual cells from three different roots per treatment were measured for fluorescence intensity. Results were statistically analyzed (ANOVA with Dunnett’s test) using SigmaPlot (San Jose, CA, USA), with significance at *P* <0.001.

### Cell cycle analysis

To assess the frequency of cells in various cell cycle stages, control and treated specimens prepared for F-actin labeling or tubulin immunostaining and DAPI staining were examined under a Zeiss AxioImager.Z2 light microscope equipped with epifluorescence or a Zeiss Observer.Z1 (Carl Zeiss AG, Munich, Germany) microscope with CLSM module. Cells at different cell cycle stages (interphase, preprophase/prophase, metaphase/anaphase and cytokinesis) were recognized according to F-actin/microtubule arrangement and/or chromatin state. At least 1000 individual cells from three different roots per treatment (a minimum of 300 meristematic cells per root) were counted. Statistical analysis of data (chi-squared test, df = 3) was performed with SigmaPlot, with significance at *P* <0.001.

### Live imaging of cytoplasmic streaming

For cytoplasmic streaming recording, 4- to 5-day-old rice seedlings were placed on glass slides and their roots were dipped either in water (control) or in the extract, covered with a coverslip and observed with DIC optics under a Zeiss AxioImager.Z2 light microscope (Carl Zeiss AG), equipped with an AxioCam MRc5 camera (Carl Zeiss AG). Time lapse images were captured using AxioVision Rel. 4.8.2 software (Carl Zeiss AG). Specifically, for each recording, 60 images captured every 2 sec, were combined in videos of 12 sec duration. The extract used for treatment was replenished regularly after each image capture.

### Transmission Electron Microscopy (TEM)

Control and extract-treated (for 30 min or 1 h) root tips of 4- to 5-day-old rice seedlings were fixed in 3% (v/v) glutaraldehyde (PolySciences, Niles, IL, USA) in 50 mM sodium cacodylate buffer (pH 7) for 4 h, washed and post-fixed with 1% (w/v) osmium tetroxide in the same buffer for 3 h at 4°C in the dark. Dehydration was performed in an acetone series, followed by treatment with propylene oxide (SERVA, Heidelberg, Germany) at 4°C and embedding in Spurr’s resin (PolySciences, Niles, IL, USA). The embedded specimens were sectioned with a Reichert-Jung Ultracut E (Reichert-Jung Optical Company, Vienna, Austria) ultramicrotome. Ultrathin sections (70-90 nm) were collected on formvar-coated copper grids and double-stained in the dark with 2% (w/v) uranyl acetate in 70% (v/v) ethanol for 15 min and 1% (w/v) lead citrate for 10 min. The sections were examined with a JEOL JEM 1011 (JEOL Ltd., Tokyo, Japan) TEM, equipped with a Gatan ES500W (Gatan Inc., Pleasanton, CA, USA) digital camera and images were acquired using Digital Micrograph 3.11.2 software. CLSM and TEM images were processed with Adobe Photoshop CS4 with only linear settings.

## RESULTS

### Effects on F-actin

Fine actin filaments were abundant in all untreated meristematic cells (Fig. 1A1). F-actin phragmoplasts were obvious in cytokinetic cells, either control or treated with MC-LR for various times (arrows in Fig. 1). In MC-LR-treated cells, cortical actin filaments appeared disoriented (indicative arrowheads in Fig. 1B2) and bundled (indicative arrowheads in Figs 1C2, D2), from 30 min until 2 h of exposure, compared to the control (Fig. 1A2). Especially after 30 min and 1 h, ring-shaped F-actin conformations (typical examples in Fig. 1C4) could be observed in affected meristematic cells (arrowheads in Figs 1B3, C3, C5). However, after 24 h of treatment, F-actin intensity deteriorated significantly (Fig. 1E1) and cortical actin filaments were barely visible (Fig. 1E2).

**Figure 1.**
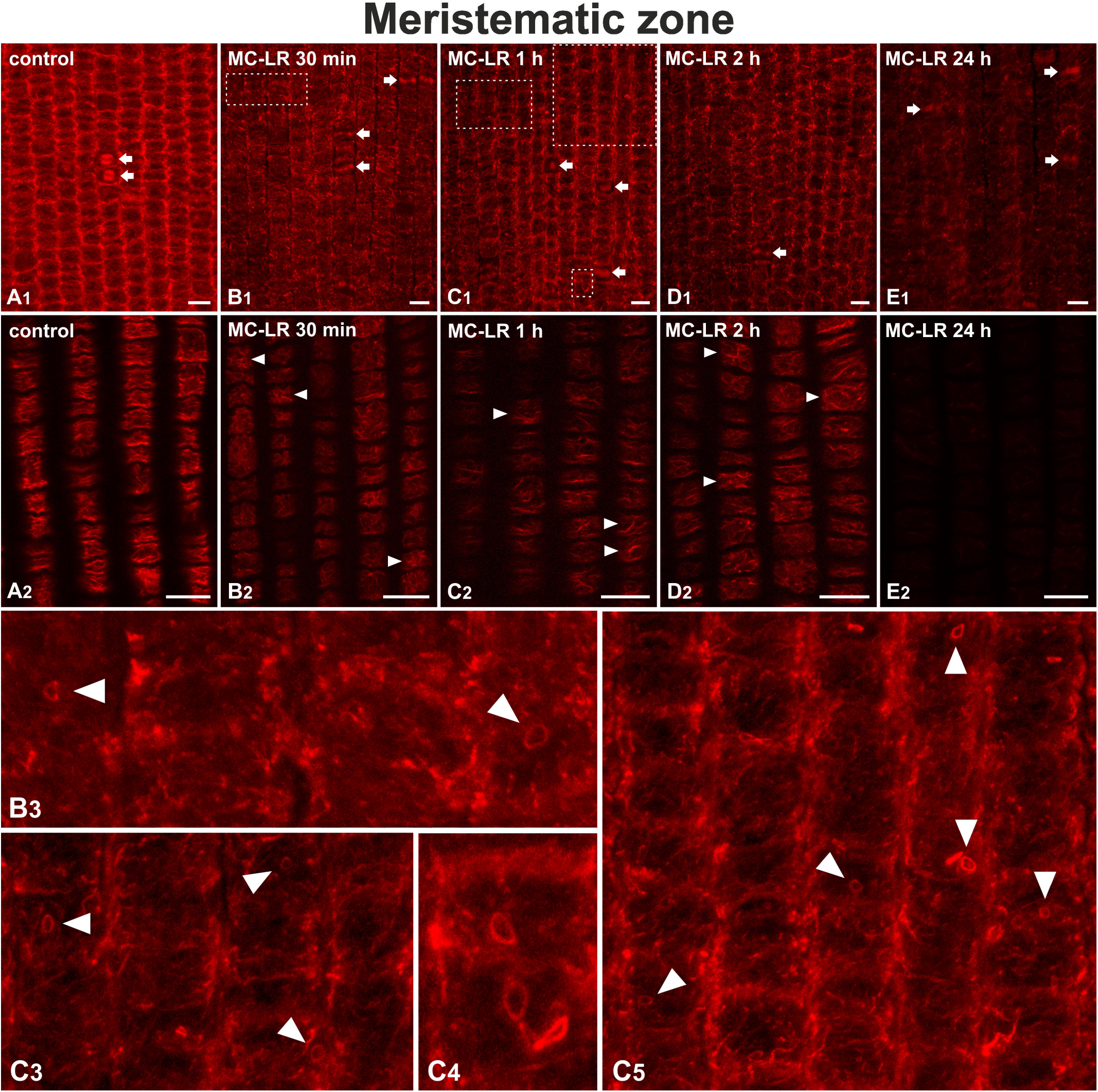
Maximum intensity projections of serial CLSM sections (A1, B1, C1, D1, E1, B3, C3-C5) and single cortical CLSM sections (A2, B2, C2, D2, E2) of *O. sativa* protodermal cells in the root meristematic zone, after F-actin staining. treatment with purified MC-LR and/or. In all images the root tip points towards the bottom of the page. Single cortical sections depict cells from the same root areas as the projections, at higher magnification. Control cells exhibit a well-organized network of abundant fine cortical and endoplasmic actin filaments (**A1**). The dominant orientation of cortical actin filaments is transverse (perpendicular to the root axis, **A2**). Cytokinetic cells with prominent F-actin phragmoplasts are noted by arrows (projections **A1**-**E1**). After treatment with MC-LR, cortical actin filaments appear disoriented at 30 min (indicative arrowheads in **B2**; *cf.* **A2**), tending to form bundles after 1 h (indicative arrowheads in **B2, C2, D2**). Ring-shaped F-actin conformations (actin-rings) are visible in affected cells after 30 min and 1 h of treatment (arrowheads in **B3** and **C3-C5**, enlargements of framed areas in **B1** and **C1**, respectively). After 24 h, F-actin fluorescence intensity is noticeably decreased (**E1**) and the cortical F-actin network barely visible (**E2**). Scale bars: 10 μm.

Meristematic cells treated with the cyanobacterial extract (Fig. 2) also exhibited disoriented cortical actin filaments after 30 min of exposure (Fig. 2A2), as well as bundling effects (indicative arrowheads in Figs 2B1, B2, C1, C2) after 1 and 2 h. After 24 h, apart from F-actin bundling (arrowhead in Fig. 2D2), several cells appeared totally devoid of actin filaments (star in Fig. 2D1).

**Figure 2.**
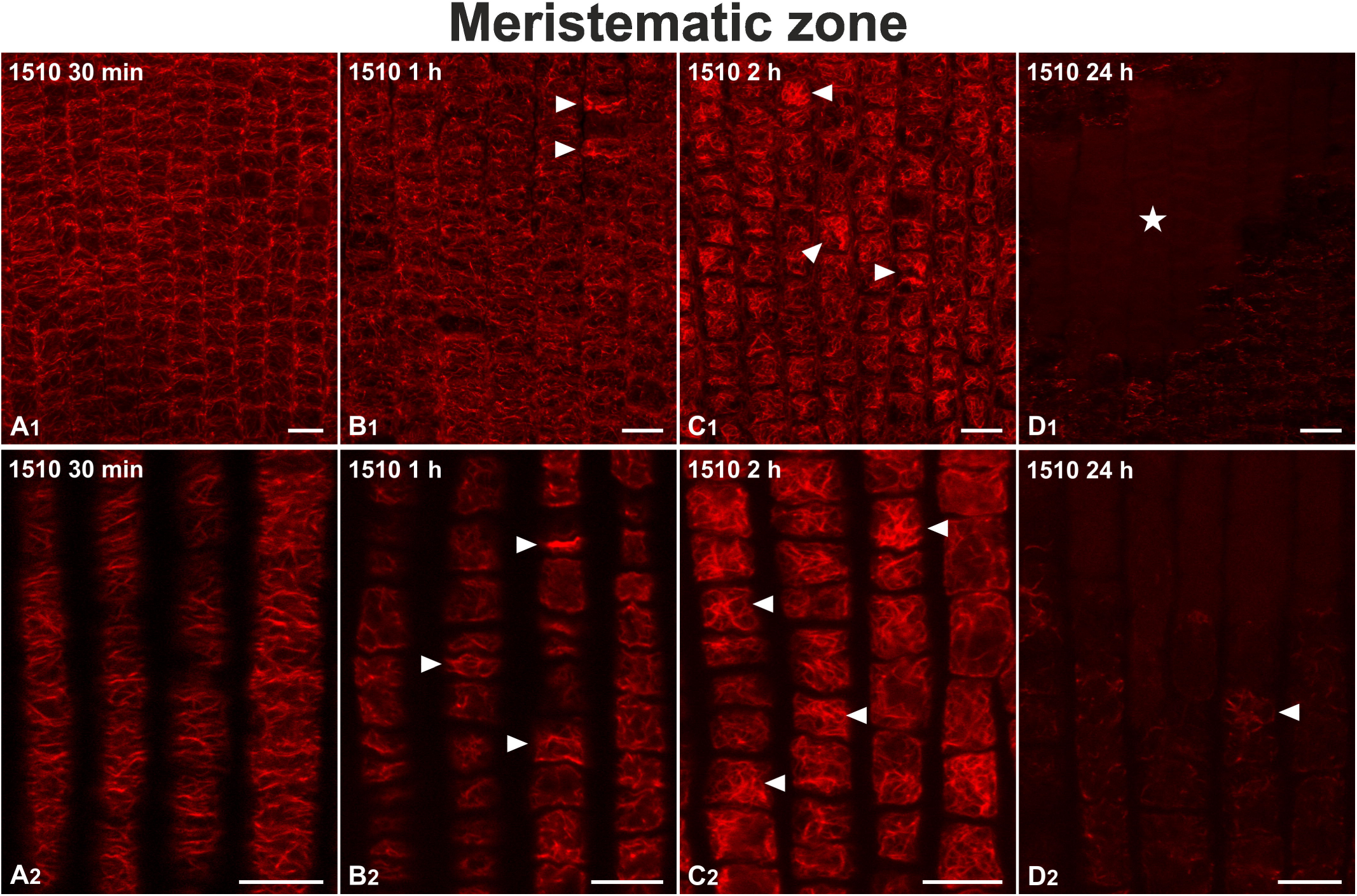
Maximum intensity projections of serial CLSM cortical sections (A1, B1, C1, D1) and single cortical CLSM sections (A2, B2, C2, D2) of *O. sativa* protodermal cells in the root meristematic zone, after treatment with the *M. flos-aquae* TAU-MAC 1510 extract and F-actin staining. All figures are oriented with the root tip to the bottom of the page. Single cortical sections (**A2**-**D2**) depict cells from the same root areas as the projections (**A1**-**D1**), at higher magnification. After 30 min of treatment with the extract (**AS1**), cortical actin filaments appear disoriented (**A2**; *cf.* **Fig. 1A2**), tending to form bundles after 1 h (indicative arrowheads in **B1, B2**). After 2 h, bundling increases (indicative arrowheads in **C1, C2**). After 24 h, the F-actin network has collapsed and cells devoid of actin filaments (area marked with star in **D1**) can be observed, along with remnants of bundles (arrowhead in **D2**). Scale bars: 10 μm.

In the differentiation zone, F-actin cables could be observed in untreated cells, exhibiting a predominant longitudinal orientation (Fig. 3A). After treatment with MC-LR, disorientation and bundling of F-actin was detectable (arrowheads in Figs 3B-E) between 30 min and 2 h of exposure, eventually leading to disappearance of F-actin in large areas of the tissue after 24 h (Fig. 3E).

**Figure 3.**
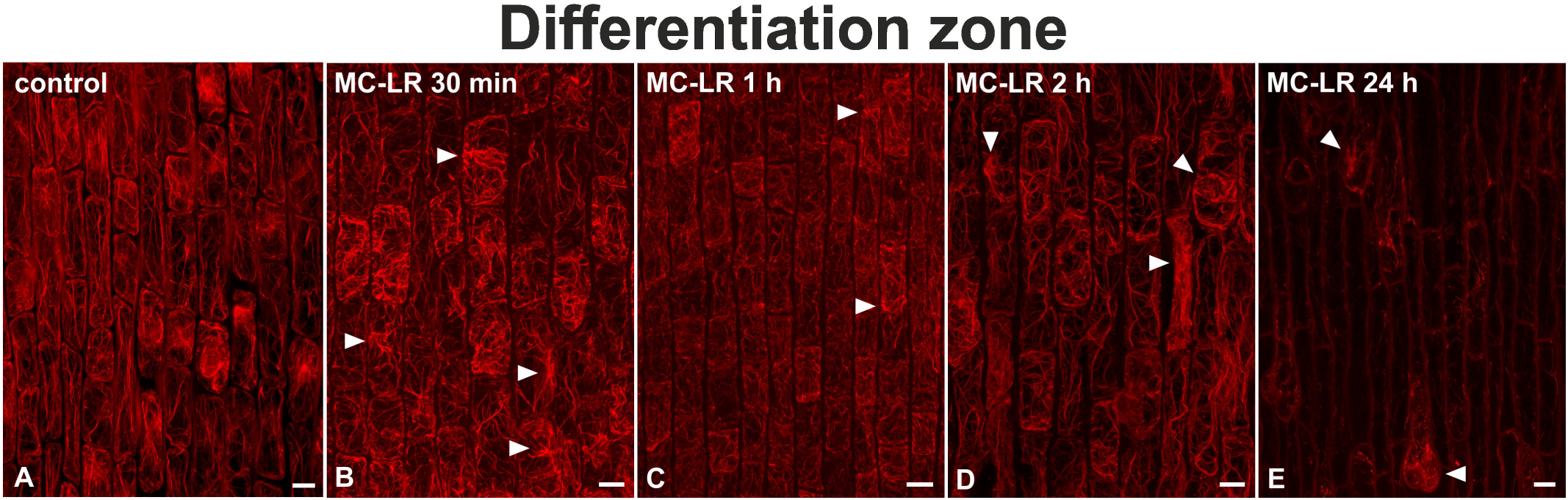
Maximum intensity projections of serial CLSM sections of *O. sativa* epidermal cells in the root differentiation zone, after F-actin staining. All figures are oriented with the root rip to the bottom of the page. In control cells, longitudinal subcortical F-actin bundles can be observed (**A**). Cells treated with MC-LR for 30 min exhibit disoriented subcortical F-actin bundles, converging at several cell sites (indicative arrowheads in **B**; *cf.* **A**). This effect is also detectable after 1 and 2 h (arrowheads in **C** and **D**, respectively). After 24 h, F-actin bundles have diminished and only remnants of F-actin bundles (arrowheads in **E**) can be observed. Scale bars: 10 μm.

In roots treated with the extract (Fig. 4), F-actin bundles were disoriented, an effect increasing in a time-dependent manner (arrows in Figs 4A, B, C1, D1). Furthermore, F-actin rings occurred in the affected cells after longer exposure (Figs 4C2, D2, D3). After 24 h, several cells were devoid of F-actin (Fig. 4D1).

**Figure 4.**
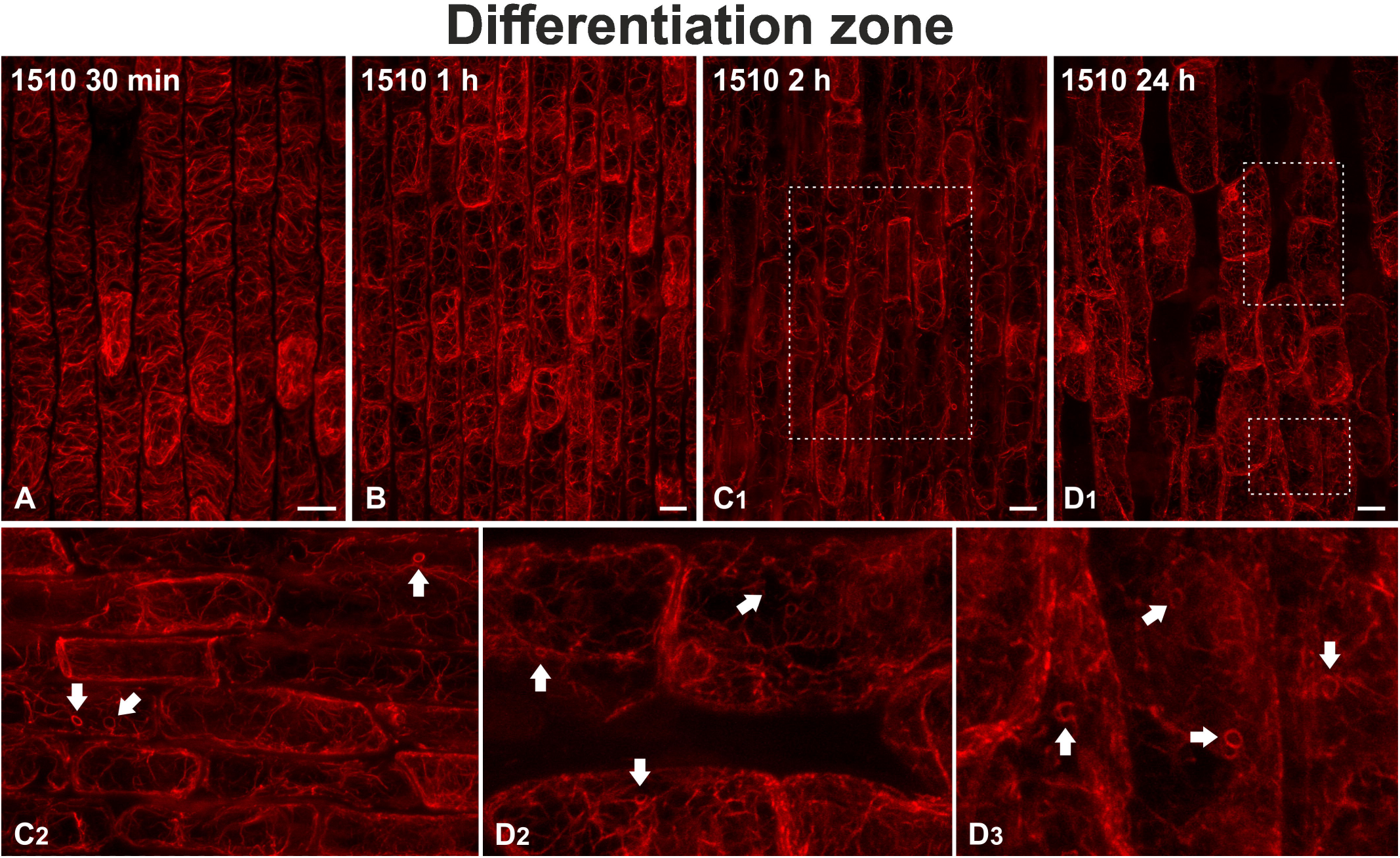
Maximum intensity projections of serial CLSM sections of *O. sativa* epidermal cells in the root differentiation zone, after treatment with the *M. flos-aquae* TAU-MAC 1510 extract and F-actin staining. All figures are oriented with the root rip to the bottom of the page, except for (**C2, D2**) with the root tip to the right. After treatment with the extract for 30 min, subcortical F-actin bundles appear to be disoriented (**A**; *cf.* **Fig. 3A**), tending to form aggregates at 1 h (**B**). After 2 h, actin filaments begin to diminish (**C1**) and F-actin rings can be observed (arrows in **C2**, enlargement of the framed area of **C1**). After 24 h, cells devoid of F-actin can be observed (**D1**), while F-actin rings (arrows in **D2** and **D3**, enlargements of framed areas of **D1**) are abundant in cells that still exhibit actin filaments. Scale bars: 10 μm.

CTCF measurements (Fig. 5) confirmed the hypothesis that fluorescence intensity gradually diminishes during the treatment with either MC-LR or the extract, in both meristematic and elongation zone the lowest levels being recorded after 24 h of treatment. This decrease appeared to be especially steep in MC-LR treated roots, just after 30 min of treatment. Any temporary increases in fluorescence intensity (e.g. in differentiated root cells after 1 h of treatment with the extract) could be attributed to F-actin bundling effects.

**Figure 5.**
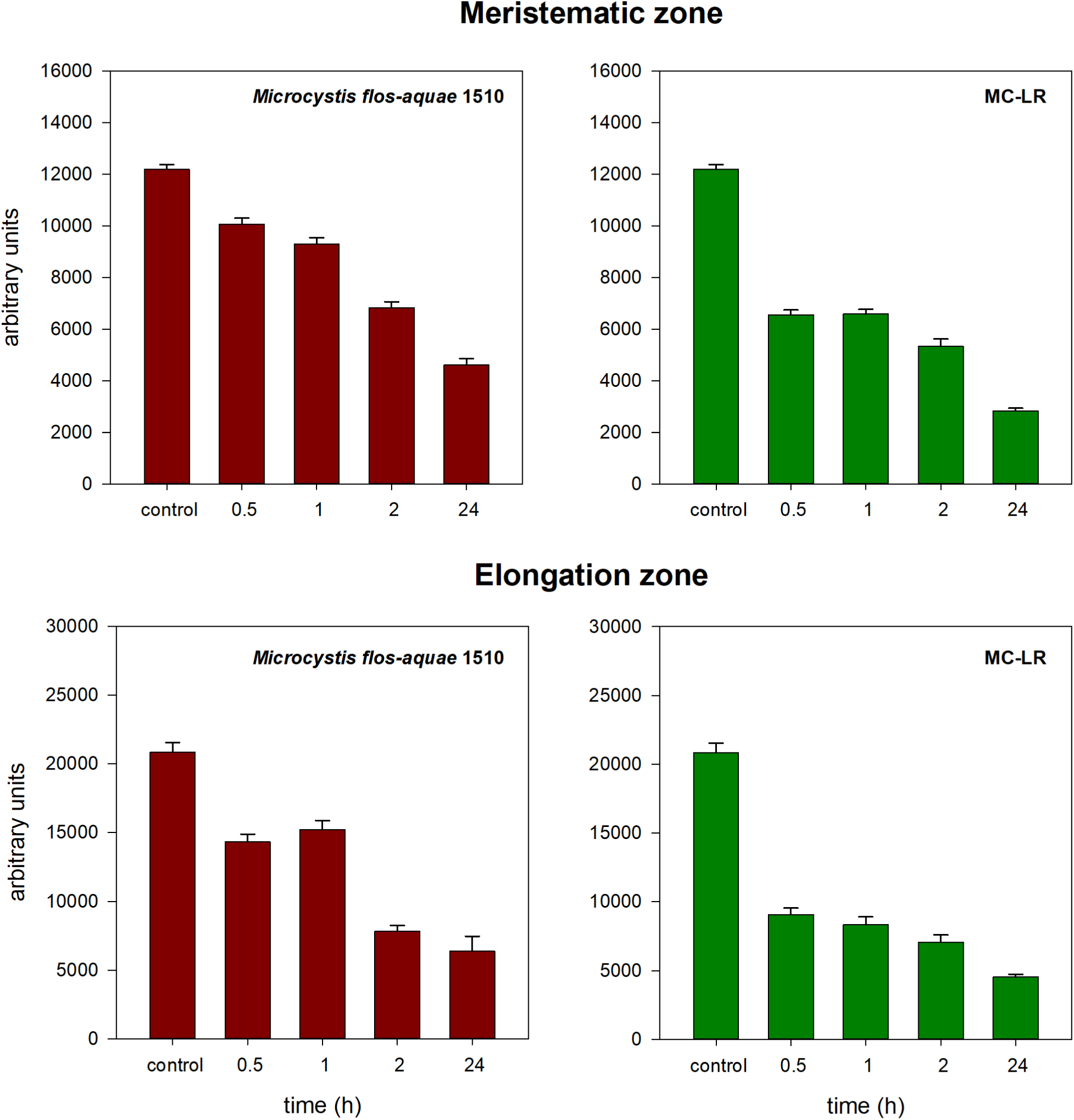
Fluorescence intensity measurements of F-actin in *O. sativa* root cells. Maximum intensity projections of serial CLSM sections of the meristematic and differentiation zones of roots treated with the *M. flos-aquae* TAU-MAC 1510 extract (maroon) or purified MC-LR (green) for various durations (30 min, 1, 2 and 24 h) were compared to similar images of control roots. Fluorescence intensity decreases drastically in a time-dependent manner, especially after treatment with MC-LR. Temporary increases, where visible, could be attributed to F-actin bundling effects. Error bars indicate the standard error. All data shown exhibit statistically significant difference compared to control (ANOVA with Dunnett’s test), *P* <0.001. *n* = 30.

### Effects on microtubules and chromatin

Untreated meristematic root cells exhibited the typical microtubule arrays, i.e. cortical microtubules in interphase cells (Fig. S1A), the preprophase band and perinuclear microtubules in preprophase/prophase cells (Fig. S1B), the mitotic spindle in metaphase/anaphase cells (Fig. S1C) and the phragmoplast in telophase/cytokinetic cells (Fig. S1D).

Rice root cells treated with the toxic cyanobacterial extract for 30 min-24 h exhibited various time-dependent alterations in the organization of microtubules (Fig. S1E-K). After 30 min of exposure, cortical microtubules of interphase cells were significantly fewer (Fig. S1E; *cf*. S1A). In affected preprophase cells, preprophase bands were visible, but perinuclear microtubules were absent (Fig. S1F; *cf*. S1B). In affected cells with condensed chromosomes, as assessed by DAPI staining, typical spindles were not found, as microtubules connected to the chromosomes appeared either abnormally elongated (Fig. S1G) or short and disoriented (Fig. S1H). In the above affected mitotic cells, microtubule fragments and/or fluorescent tubulin structures, apart from chromosome-connected microtubules, could also be observed (Figs S1G, H). In telophase/cytokinetic cells treated as above, phragmoplasts could be observed, the microtubules of which were longer than those of untreated cells (Fig. S1I; *cf*. S1D), their (-) ends sometimes attached on the surface of daughter nuclei. After 1 h of treatment, microtubules were depolymerized (Figs S1J, K) in cells with normal-looking nucleus (Fig. S1J), as well as in cells with abnormal chromatin condensation (Fig. S1K). In some of these cells, fluorescent tubulin spots could be observed (Fig. S1J).

Interestingly, MC-LR-treated root cells exhibited control-like microtubule arrays. Occasionally, 30 min- and 1 h-treated interphase cells with typical cortical microtubules (Figs S2C, E; *cf.* S2A) exhibited endoplasmic microtubules, not typically observed in untreated cells (Figs S2D, F; *cf.* S2B), while in affected preprophase cells preprophase bands exhibited gaps (Fig S2G; *cf.* Fig. S2H). MC-LR-affected metaphase cells appeared to have typical mitotic spindles (Figs S2I, J; *cf.* Fig. S1C), while occasionally chromosomes out of the spindle could be observed (arrow in Fig. S2J).

### Effects on cell cycle progression

The assessment of cell cycle stages in treated root tip cells revealed that each stage frequency was significantly altered after 1 h of treatment with MC-LR, while it was severely disturbed in extract-treated roots even after 30 min (Table 1). In the latter case, the percentage of preprophase/prophase cells was increased after 30 min of treatment, as was the percentage of metaphase/anaphase cells after 1 h, compared to untreated roots. On the contrary, there was a notable decrease in the percentage of cytokinetic cells after 30 min and 1 h of treatment with the extract. Alterations observed (in both MC-LR- and extract-treated roots) were statistically significant (chi-squared test, df = 3, *P* <0.001)

**Table 1.** Occurrence of various cell cycle stages in *O. sativa* root tips, either control or after treatment with microcystin-LR (MC-LR) or the *Microcystis flos-aquae* TAU-MAC 1510 extract for 30 min and 1 h. Data represent absolute cell counts (percentages in parentheses). Data exhibiting statistically significant difference compared to the control are noted with asterisks (chi-squared test, df = 3, *P* <0.001).

|  |  | MC-LR |  | *Microcystis flos-aquae* 1510 |  |
| --- | --- | --- | --- | --- | --- |
|  | Control (%) | 30 min (%) | 1 h (%)* | 30 min (%)* | 1 h (%)* |
| Interphase | 1080 (91.6) | 1012 (90.68) | 978 (92.44) | 895 (89.5) | 914 (91.4) |
| Preprophase/Prophase | 44 (3.73) | 53 (4.75) | 44 (4.16) | 79 (7.9) | 51 (5.1) |
| Metaphase/Anaphase | 12 (1.02) | 8 (0.72) | 7 (0.66) | 12 (1.2) | 31 (3.1) |
| Cytokinesis | 43 (3.65) | 43 (3.85) | 29 (2.74) | 14 (1.4) | 4 (0.4) |

### Effects on cytoplasmic streaming

Alterations of cytoplasmic streaming in affected root cells were visible after treatment with the extract (see Supplementary Videos). After 1 h, streaming either stopped or appeared to be slower (left and right arrow, respectively, Video S5; *cf.* S4), as opposed to that of root cells in presence of water, in which streaming remained vivid during the exposure time period (Videos S1-3). After 2 h, streaming (where present) was noticeably abnormal and cytoplasmic aggregates could be observed (arrow and arrowhead, respectively, Video S6; *cf.* S4).

### Effects on the endoplasmic reticulum and Golgi apparatus

The toxic extract also affected the integrity and distribution of the endoplasmic reticulum. In cells exposed to the extract for 30 min and 1 h, fluorescent endoplasmic reticulum aggregates were observed by CLSM, located cortically and/or around the nucleus (Fig. 6B, C), in contrast to the evenly distributed endoplasmic reticulum of untreated cells (Fig. 6A). After 3 h of treatment, these aggregates appeared to fade, while the nucleus exhibited morphological alteration (Fig. 6D).

**Figure 6.**
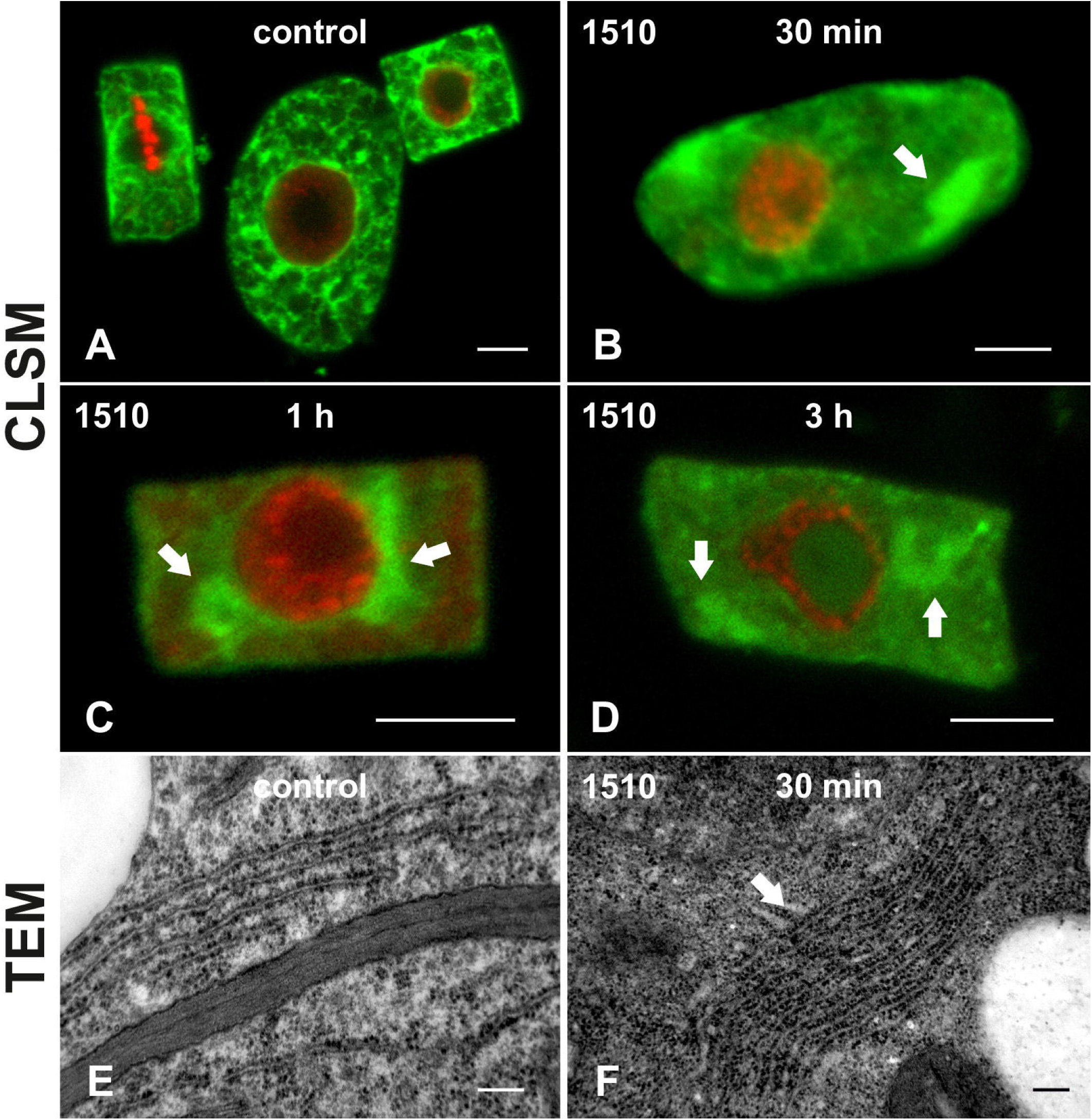
Endoplasmic reticulum distribution in *O. sativa* root cells. Various durations of treatment with the *M. flos-aquae* TAU-MAC 1510 extract are indicated on the images. **A**-**D**. Single CLSM central sections, after endoplasmic reticulum immunolabeling (green) and DNA staining with DAPI (pseudocoloration in red). Untreated cells (**A**) exhibit even endoplasmic reticulum distribution during interphase (cells at middle and right), while during metaphase endoplasmic reticulum encages the spindle (left). Treatment with the extract results in appearance of endoplasmic reticulum aggregates (arrows in **B-D**), even after only 30 min (**B**), the fluorescence intensity of which fades as treatment duration increases (**C, D**). Scale bars: 5 μm. **E, F**. TEM micrographs depicting normal endoplasmic reticulum distribution in control cells (**E**) and a stack of endoplasmic reticulum cisternae (arrow in **F**) after 30 min of treatment. After longer treatment, endoplasmic reticulum cisternae could not be found by TEM. Scale bars: 0.2 μm.

TEM observations of root cells treated with the extract for 30 min revealed that endoplasmic reticulum cisternae were heavily stacked (Fig. 6F), in contrast to the loosely packed cisternae of untreated cells (Fig. 6E). In addition, while in untreated cells Golgi dictyosomes were evenly distributed (Fig. 7A), in cells affected by the extract the dictyosomes appeared clustered, even after 30 min of treatment with vesicles trapped among them (Fig. 7B). After 1 h of exposure, Golgi apparatus clustering and vesicle aggregation were intensified (Fig. 7C).

**Figure 7.**
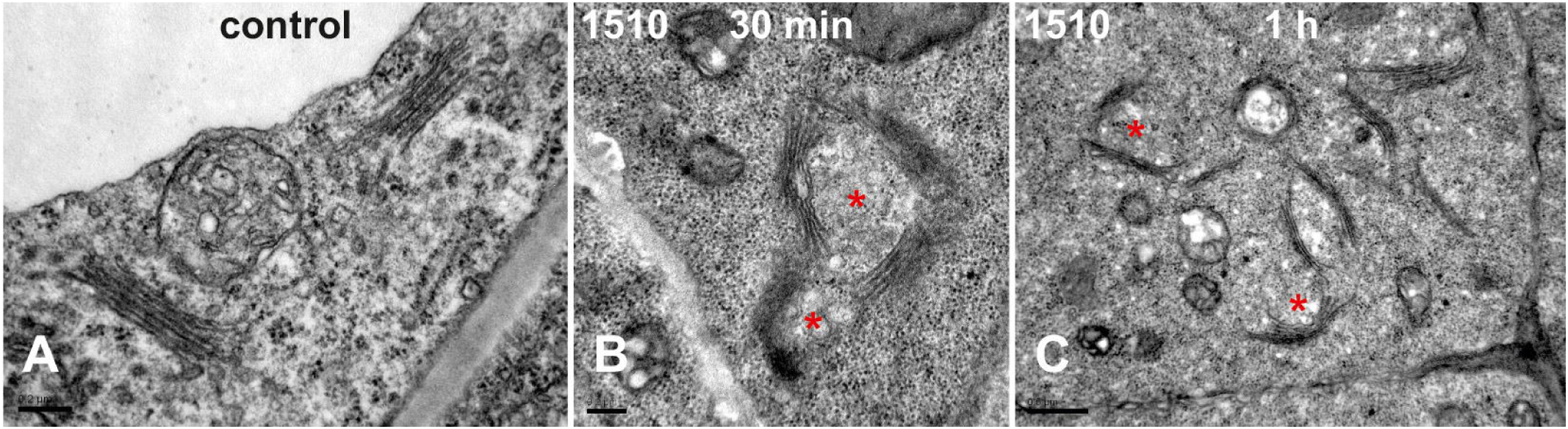
TEM micrographs of Golgi dictyosome distribution in *O. sativa* root cells. Dictyosomes are normally distributed and distanced in an untreated cell (**A**). After 30 min (**B**) and 1 h (**C**) of treatment with the *M. flos-aquae* TAU-MAC 1510 extract, dictyosomes gather in clusters, displaying aggregates of vesicles (asterisks) entrapped between them. Scale bars: 0.2 μm (**A, B**) and 0.5 μm (**C**).

## DISCUSSION

In this study, MC-LR disrupted F-actin in rice roots. The detrimental effects were recorded as significant alterations of the network (Figs 1, 3) and as a gradual decrease in F-actin fluorescence intensity (Fig. 5), eventually leading to a collapse in both meristematic and differentiating root cells (Figs 1, 3).

Apart from the above similarity, there was a significant difference between the treatment with purified MC-LR and the extract. Exposure to the latter induced severe disruption of microtubule arrays and alterations of the frequency of cells at various cell cycle stages after just 30 min (Table 1). On the contrary, MC-LR affected microtubule organization only slightly, as well as cell cycle distribution only after 1 h of exposure.

According to cytological studies on other plant species, MCs affect the integrity and organization of microtubules, causing chromatin hypercondensation in plant cells by inhibiting PP1 and PP2A, which subsequently leads to histone H3 hyperphosphorylation (Beyer et al., 2012; Máthé et al., 2009; Ujvárosi et al., 2019). Also, experiments on *Arabidopsis thaliana* indicated the involvement of subfamily II and TON2 subunits of the PP2A holoenzyme in cortical microtubule organization and, probably, in microtubule nucleation through interaction with γTuRCs (Yoon et al., 2018). Studies on PP2A have highlighted its importance for plant cytoskeleton organization (especially microtubules), affecting preprophase band formation, division plane determination (Spinner et al., 2013), and cell shape (Kirik et al., 2012). Interestingly, in the present study, treatment with purified MC-LR did not result in any of these effects. A possible reason for this could be that the duration of treatments here was far shorter than in the previously mentioned studies. In most of the relevant studies, exposure to MC-LR spanned from 4 h to 20 days (Beyer et al., 2012; Garda et al., 2016; Mathé et al., 2009). However, in the present study microtubules were severely affected only after treatment with the crude toxic extract for a very short time (30 min and 1 h), which indicates the possible involvement of other compounds in the effect mechanism. Except of microcystins, which have been detected in *Microcystis flos-aquae* TAU-MAC 1510, it is also likely that these effects may be the result of a synergistic action of various bioactive compounds produced by the strain. For example, anabaenopeptins, which show activity against PP1 (Gkelis et al., 2006), are known to occur in large concentrations in *Microcystis*-dominated blooms (Gkelis et al., 2015b) or microginins, which have been recently reported to be abundant in blooms of Greek freshwater cyanobacteria (Zervou et al., 2020), are also known to be bioactive (Ujvárosi et al., 2020; for a review on the variety of cyanobacterial bioactive compounds, see Dittmann et al., 2015; Elisabeth & Janssen, 2019).

The findings of the present study prove that F-actin is indeed a target of cyanobacterial toxicity, specifically of microcystins, in plant cells. To date, information about the way actin filaments are affected by microcystins derives exclusively from studies on treated animal cells. In rat hepatocytes and fibroblasts, all three cytoskeletal components (microtubules, intermediate filaments and microfilaments) are sensitive to MC-LR treatments in a dose- and time-dependent manner (Wickstrom et al., 1995). Disorganization of intermediate filaments and microtubules was reported to precede alterations in normal disposition of microfilaments in animal cells. These alterations included extensive reorganization of F-actin into “rosette-like structures”, which eventually collapsed, forming aggregates around the cell nucleus (Wickstrom et al. 1995). Similar observations concerning F-actin were also made in rat hepatocytes by Ghosh et al. (1995) and by Batista et al. (2003) in primary human hepatocytes, while Gácsi et al. (2009) reported the formation of circular actin aggregations around the nuclei of MC-LR-treated Chinese hamster ovary (CHO-K1) cells after 24 h of incubation at high concentrations of the cyanotoxin (20 μM). In particular, actin aggregation at the cell periphery seems to be a common alteration in microcystin-treated animal cells (Eriksson et al., 1989; Falconer and Yeung, 1992; Runnegar and Falconer, 1986; Wickstrom et al., 1995), at least for some time during the cascade of F-actin-collapse events. As expected, alterations in F-actin organization consequently led to animal cell shape deformation.

It could be suggested that F-actin disruption in rice root cells is attributed to protein phosphatase inhibition. In human T lymphocytes, inhibition of PP1 and PP2A has been shown to block dephosphorylation of actin depolymerizing factor (ADF)/cofilin, an actin-binding protein, which is responsible for actin depolymerization when in active, unphosphorylated state (Ambach et al., 2000). Such an effect on PP1 and PP2A could explain the bundling effects observed in the F-actin network of rice root cells affected by microcystin, as F-actin tends to become stabilized. In a more recent study (Wang et al., 2014), MC-LR was found to directly bind to PP2A in SMMC-7721 human liver cancer cells, inhibiting enzyme activity and causing cytoskeletal rearrangements. More specifically, PP2A inhibition led to hyperphosphorylation of various cytoskeleton-associated proteins (including cofilins) and inactivation (as well as changes in subcellular localization) of Rac1, a small GTPase, which is regulated by PP2A (Nunbhakdi-Craig et al., 2003) and is involved in microtubule and actin dynamics (Wittmann et al., 2003). In plants, the presence of cofilins has been reported in various species (Hussey et al., 2002), while PP2A-2, an isoform of PP2A, is known to directly regulate ADF/cofilin (and, therefore, F-actin rearrangements) in *Arabidopsis thaliana* (Wen et al., 2012). In addition, OsRac1 (a rice homolog of Rac1), along with its activator GEF protein OsSPK1, have been recently suggested to be regulators of actin dynamics in rice (Wang et al., 2018).

Importantly, in rice root cells, progression of F-actin disorganization/disorientation appeared to be slower than in animal cells and not as harsh; in fact, even after 24 h of exposure, actin filaments could still be observed, though scarce. The increased resistance of actin filaments to microcystins could be attributed to at least three reasons: (1) Most of the actin filaments, especially in vacuolated cells, are interconnected in thick bundles, which could offer stabilization against adverse factors. (2) Cortical actin filaments are interconnected to cellulose microfibrils of the cell wall by formin1 (Martinière et al., 2011), also regulating F-actin stability and cell shape (Rosero et al. 2013, 2016). (3) F-actin may appear more stabilized following inactivation of ADF/cofilin, due to inhibition of protein phosphatases by microcystins. Whichever the mechanism by which F-actin is affected by microcystins, the effect of a microcystin-rich crude extract indicates that this could occur in environmental conditions, due to cell lysis during cyanobacterial blooms. According to this view, disruption of F-actin may be one more possible reason for cyanobacterial toxicity to plants, apart from the already established effects on microtubules and the cell cycle. In addition, the importance of F-actin disorganization is also reflected on the alterations observed in endoplasmic reticulum and the Golgi apparatus. As reviewed by Volkmann and Baluška (1999), movement and distribution of endoplasmic reticulum and the Golgi apparatus in plant cells are controlled by the F-actin network. Furthermore, cytoplasmic streaming was disturbed (see Supplementary Videos), due to its dependence on actomyosin integrity and function (for a review, see Shimmen and Yokota, 2004). Consequently, stacking of endoplasmic reticulum as well as abnormal aggregation of Golgi dictyosomes could be a direct result of defective F-actin organization and subsequent intracellular motility defects. Apart from the involvement of F-actin, alterations of endoplasmic reticulum have been reported in both plant (Huang et al., 2009) and animal cells, either fish (Li et al., 2001) or mammalian (Alverca et al., 2009), and attributed to microcystins, after treatment for various time periods, ranging from hours to days. These results, therefore, suggest the presence of a mechanism of toxicity, which involves microcystins and affects a variety of cell systems. Nevertheless, in the present study, endoplasmic reticulum cisternae seemed to diminish after 1 h of exposure. Interestingly, absence of smooth endoplasmic reticulum has been also previously observed in rat hepatocytes after treatment with MC-LR for 20 min (Eriksson et al., 1989).

In conclusion, F-actin is a target of MC-LR. In addition, the microcystin-rich extract of the cyanobacterial strain *Microcystis flos-aquae* TAU-MAC 1510 disrupts cytoskeletal components, F-actin and microtubules, as well as the endomembrane system and cell cycle distribution, in root tip cells of *Oryza sativa*. Microcystins present in the extract could be held responsible for certain of the effects observed, as they have previously been linked with defects in chromatin condensation, but most probably a synergistic action of microcystins with other cyanobacterial bioactive compounds may be hypothesized. To our best knowledge, this is the first report of a microcystin (MC-LR) affecting the plant F-actin cytoskeleton and the first attempt to feature the potent effects of cyanobacterial bioactive compounds on F-actin-related cell functions. Further research is needed in order to elucidate the above in detail.

## ACKNOWLEDGEMENTS

This research is part of a PhD thesis (Dimitris Pappas) and is co-financed by Greece and the European Union (European Social Fund-ESF) through the Operational Programme «Human Resources Development, Education and Lifelong Learning» in the context of the project “Strengthening Human Resources Research Potential via Doctorate Research” (MIS-5000432), implemented by the State Scholarships Foundation (IKY). Emmanuel Panteris is supported by the AUTh Research Committee (Grant No 91913). The authors would like to deeply thank Prof. Kaarina Sivonen (University of Helsinki, Finland) for generously providing a stock of MC-LR, as well as Assist. Prof. Ioannis-Dimosthenis Adamakis (National and Kapodistrian University of Athens, Greece) and Assoc. Prof. George Komis (Palacký University Olomouc, Czechia) for their advice and critical reading of the manuscript.

