## SUPPLEMENTARY DATA LEGENDS for "The effects of microcystin-LR in *Oryza sativa* root cells: F-actin as a new target of cyanobacterial toxicity"

**Figure S1. Maximum intensity projections of serial CLSM sections (A-C, E-H, J, K) and central single CLSM sections (D, I) of *O. sativa* root meristematic cells, after *α*-tubulin immunostaining (green) and DNA staining with DAPI (pseudocoloration in red).** The duration of exposure to the *M. flos-aquae* TAU-MAC 1510 extract is indicated on each figure. **A**-**D**. Control cells. At interphase (**A**), a dense network of transverse cortical microtubules is present. In preprophase/prophase cell (**B**; see chromatin condensation) a preprophase microtubule band (defined by brackets) can be observed, while perinuclear microtubules converge to two prominent poles (pointed by arrows). At metaphase (**C**), a typical spindle with chromosomes aligned at its equator is present, while cell at cytokinesis (**D**) exhibits a typical phragmoplast. After 30 min of treatment with the extract (**E**-**I**), cortical microtubules of interphase cell (**E**) appear diminished (*cf*. **A**), while at preprophase (**F**; see chromatin condensation, typical of preprophase/prophase cell) a preprophase microtubule band is present (brackets in **F**), but no perinuclear microtubules can be observed (*cf*. **B**). In mitotic cells, as assessed by chromosome masses (**G**, **H**), microtubules are not organized in typical spindles, but either are elongated (**G**) or appear short and disorganized (**H**). Also note the fluorescent speckles in (**H**). At cytokinesis (**I**), a phragmoplast can be observed, the microtubules of which appear abnormally long (*cf*. **D**). **J**, **K**. After 1 h of treatment, microtubules have totally disappeared. Note the fluorescent patches, probably tubulin aggregates, in (**J**) and the highly condensed chromatin aggregate in (**K**). Scale bars: 5 μm.

**Figure S2. Cortical (A, C, E, G) and central (B, D, F, H) single CLSM sections, and maximum intensity projections of serial CLSM sections (I, J) of *O. sativa* root meristematic cells, after *α*-tubulin immunostaining (green) and DNA staining with DAPI (pseudocoloration in red).** The duration of treatment with purified MC-LR is indicated on each figure. **A**, **B**. Control interphase cells exhibit typical transverse cortical microtubules (**A**) and no endoplasmic ones (**B**). **C**-**F.** MC-LR-affected interphase cells after 30 min (**C**, **D**) and 1 h (**E**, **F**) of treatment, exhibiting apart from typical cortical microtubules (**C**, **E**; *cf.* **A**) also endoplasmic microtubules (**D**, **F**; *cf.* **B**). **G**, **H**. MC-LR-affected preprophase cell after 1 h of treatment, exhibiting a preprophase microtubule band at cortical level (**G**), but not at lateral cell faces at central section (arrows in **H** point to the sites of expected preprophase band presence; *cf.* brackets in **Fig. S1B**). **I**, **J**. Metaphase cells after 30 min (**I**) and 1 h (**J**) of treatment exhibit typical mitotic spindles (*cf.* **Fig. S1C**). Note the chromosome outside the spindle, observed after 1 h of treatment (arrow in **J**). Scale bars: 5 μm.

**Video S1.** Cytoplasmic streaming of *O. sativa* root cells at the beginning of exposure to water (t=0). Scale bar: 10 μm.

**Video S2.** Cytoplasmic streaming of *O. sativa* root cells after 1 h of exposure to water (t=1 h). Scale bar: 10 μm.

**Video S3.** Cytoplasmic streaming of *O. sativa* root cells after 2 h of exposure to water (t=2 h). Scale bar: 10 μm.

**Video S4.** Cytoplasmic streaming of *O. sativa* root cells at the beginning of exposure to the *M. flos-aquae* TAU-MAC 1510 extract (t=0). Cells of interest are noted with arrows. Scale bar: 10 μm.

**Video S5.** Cytoplasmic streaming of *O. sativa* root cells after 1 h of exposure to the *M. flos-aquae* TAU-MAC 1510 extract (t=1 h). Cells of interest are noted with arrows. Scale bar: 10 μm.

**Video S6.** Cytoplasmic streaming of *O. sativa* root cells after 2 h of exposure to the *M. flos-aquae* TAU-MAC 1510 extract (t=2 h). Cells of interest are noted with arrows. Scale bar: 10 μm.
