## Supplementary figures and images for "The effects of microcystin-LR in *Oryza sativa* root cells: F-actin as a new target of cyanobacterial toxicity"

### Fig. S1

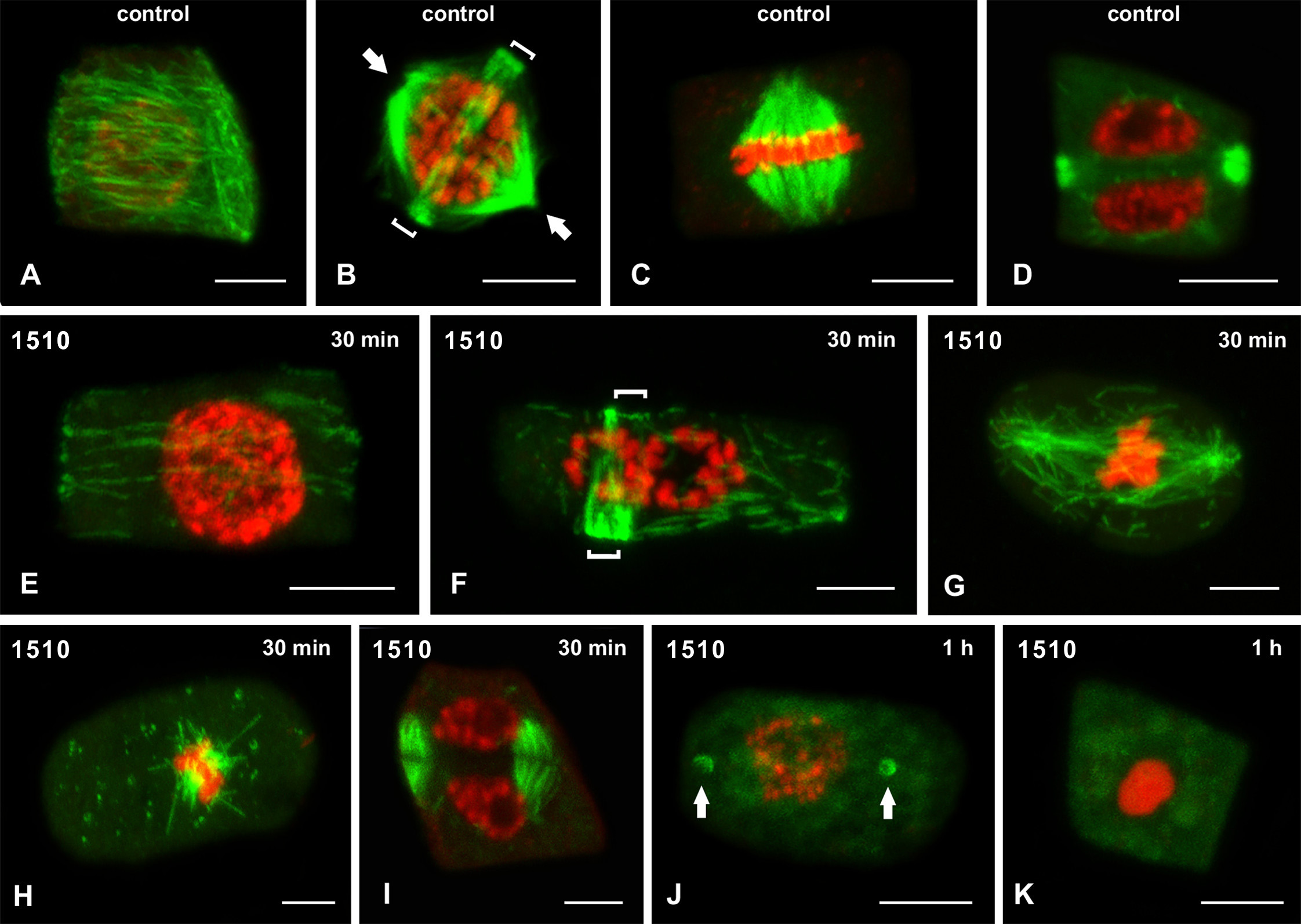

### Fig. S2

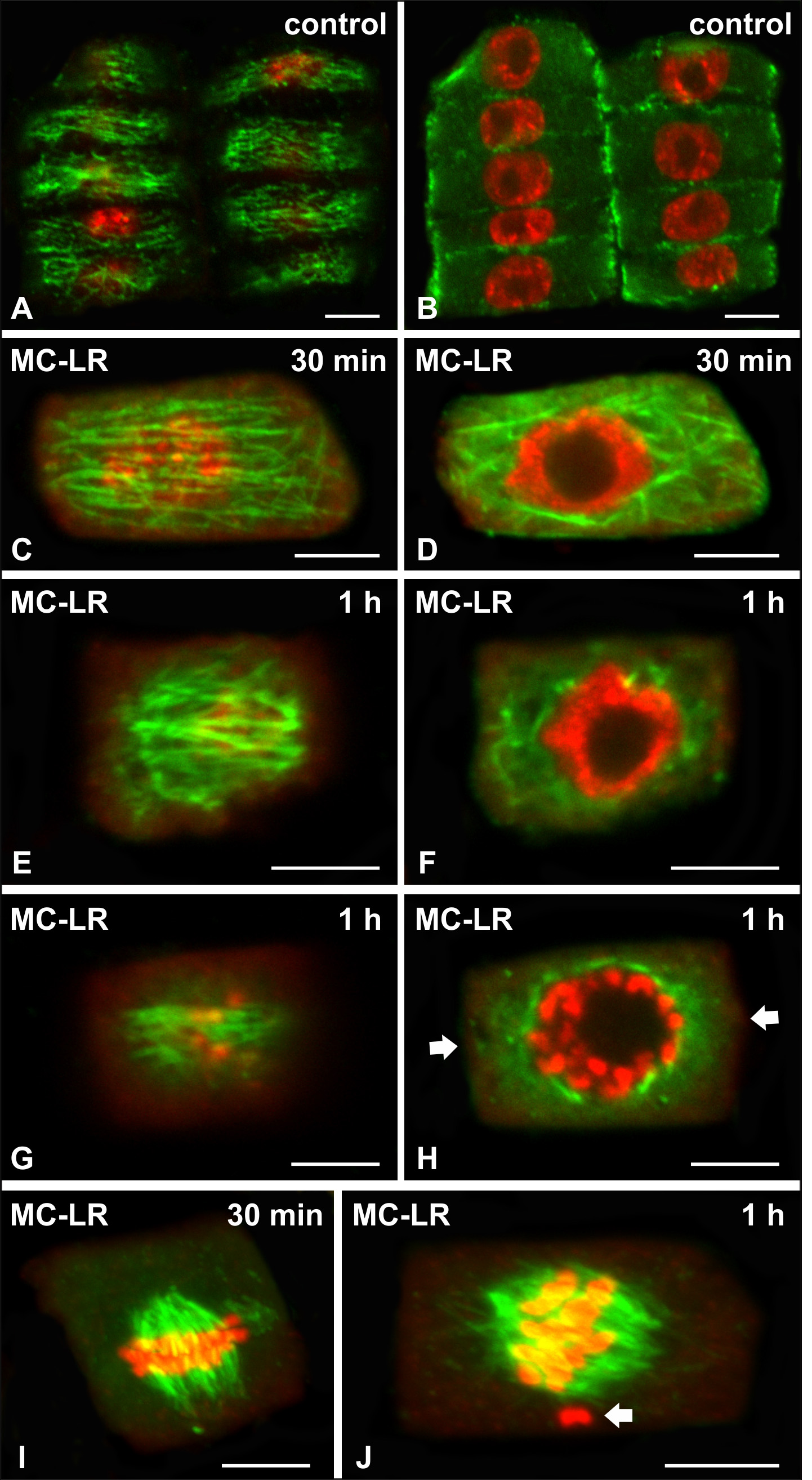
